# Dynamic Responses of Striatal Cholinergic Interneurons Control the Extinction and Updating of Goal-Directed Learning

**DOI:** 10.1101/2023.11.09.566460

**Authors:** Zhenbo Huang, Ruifeng Chen, Matthew Ho, Xueyi Xie, Xuehua Wang, Jun Wang

## Abstract

Striatal cholinergic interneurons (CINs) regulate behavioral flexibility, but their exact contribution to this process remains elusive. In this study, we report that extinction learning enhances acetylcholine (ACh) release. Mimicking this enhancement by optogenetically inducing CIN burst firing promotes extinction learning. CINs receive excitatory thalamic inputs, and we observed that extinction training augmented thalamic activity. Optogenetically stimulating these thalamic inputs caused CIN burst firing and enhanced ACh release, strengthening extinction learning. Notably, CIN burst firing is usually followed by a pause in firing. We found that disrupting this pause through continuous optogenetic stimulation reversibly impaired the updating of goal-directed behaviors. Furthermore, excessive alcohol consumption reduced thalamus-induced burst-pause firing in CINs and impaired the reversal of goal-directed learning. In summary, thalamic-driven CIN burst firing promotes extinction learning, while the pause is pivotal for reversing goal-directed behavior, a process impacted by excessive alcohol. These findings shed light on how CIN’s dynamic responses affect behavioral flexibility.

**Highlights:** H1. **Burst** firing of CINs promotes extinction learning

H2. Thalamic-CIN **excitation** enhances extinction learning

H3. **Pause** of CIN is critical for the reversal of goal-directed learning

H4. Chronic alcohol consumption reduces the **burst**-**pause** of **CINs** and impairs the reversal of goal-directed learning.

## Introduction

Behavioral flexibility, the ability of an organism to adapt its behavior to changing circumstances or environmental cues, is critical for species survival [1]. Central to this capability is the basal ganglia, which orchestrates selecting and implementing contextually appropriate responses based on expected outcomes in conjunction with the prefrontal cortex [2–4]. Within the basal ganglia, the striatum, which predominantly receives cognitive inputs from the cerebral cortex and sensory information from the thalamus, stands out as an integrator facilitating voluntary action initiation, execution, and modulation [5–7]. Acetylcholine (ACh) in the dorsomedial striatum (DMS) has been implicated in behavioral flexibility [8, 9]. Further studies have identified the cholinergic interneurons (CINs) in the posterior DMS as crucial for behavioral flexibility [10–12].

Even though CINs constitute a mere 1-2% of the striatal cell population, their extensive terminal fields pervade the entire striatum [13]. CINs provide a major source of ACh to the striatum and modulate striatal output by regulating other types of neurons [14–16]. CINs were shown to correspond to putative “tonically active neurons” (TANs) in the early *in vivo* studies of primates [17]. They were initially thought to have minimal behavioral state modulation due to their consistent firing during movements [18]. Subsequent research, however, revealed their responsiveness to reward cues, characterized by a “pause response” either after a brief burst firing or before a “rebound” firing [15, 19–22]. These CIN dynamic responses to motivationally salient stimuli coincide with phasic changes in dopamine neuron activity [22]. Increasing literature suggests that acetylcholine and dopamine systems interact in reward processing within the striatum [17, 22–24]. However, the details of CIN dynamics and their significance across behavioral contexts largely remain elusive.

The mechanisms of CIN dynamic responses are not entirely clear. Different hypotheses have been proposed [25, 26]. Evidence suggests that the activation of thalamic inputs can induce a burst-pause firing pattern in the CINs [20, 27]. The intralaminar thalamic nuclei, especially the parafascicular nucleus (PfN), provide a major excitatory input to CINs [28, 29]. An *in vivo* study with anesthetized rats showed electrical stimulation of the thalamus-induced pauses in CIN firing [20]. A primate study also indicated that thalamic inputs were necessary for the pause response of TANs, as local inhibition of thalamic activity by a GABA_A_ receptor agonist, muscimol, attenuated the pauses [30]. It has been proposed that these pauses provide a time window during which corticostriatal synaptic plasticity can occur [31]. Indeed, a recent study using anesthetized rats demonstrated that the pauses in CIN firing were required to induce corticostriatal plasticity [32]. However, the behavioral implications of CIN dynamic responses are still unclear. Our study aims to bridge this knowledge gap. Using goal-directed instrumental learning, we reveal the CINs’ critical functions in extinction learning and behavioral adaptation. Specifically, while CIN burst firing accelerates the extinction of acquired behaviors, the pauses in CIN firing prove essential for updating goal-directed learning.

## Results

### Burst firing of CINs promotes extinction learning

Prior research has shown that CINs respond to salient stimuli [15, 19, 22]. As the omission of rewards could be perceived as salient stimuli by animals, we hypothesized that extinction training alters CIN activity. To test this hypothesis, we infused a genetically encoded green ACh sensor, AAV-iAChSnFR [33–35], into the DMS of Long-Evans rats (Fig. 1A) and verified its expression (Fig. 1B). The sensor displayed a dose-response to varying ACh concentrations but remained unresponsive to dopamine (Fig. 1C, D). After operant training associated lever pressing with sucrose rewards, rats participated in a combined acquisition and extinction session. Within this combined session, rats pressed the lever to receive sucrose rewards for the initial ten trials, establishing a baseline for the ACh signal. The subsequent ten trials involved lever pressing, yielding no reward, representing the extinction phase. Analysis of the signal heat map revealed that the ACh fluorescence dropped immediately following reward delivery, followed by a pronounced rise. Notably, this fluorescence rise during extinction was more intense and extended than in baseline trials (Fig. 1E, F). Quantitative assessments of peak signals and the area under the curve (AUC) revealed that extinction led to enhanced and sustained ACh release in the DMS (Peak: Fig. 1G, *t*_(5)_ = -5.13, *p* < 0.01; AUC: Fig. 1H, *t*_(5)_ = -3.19, *p* < 0.05). Our data suggests that extinction training heightened ACh signaling in the DMS.

**Figure 1.**
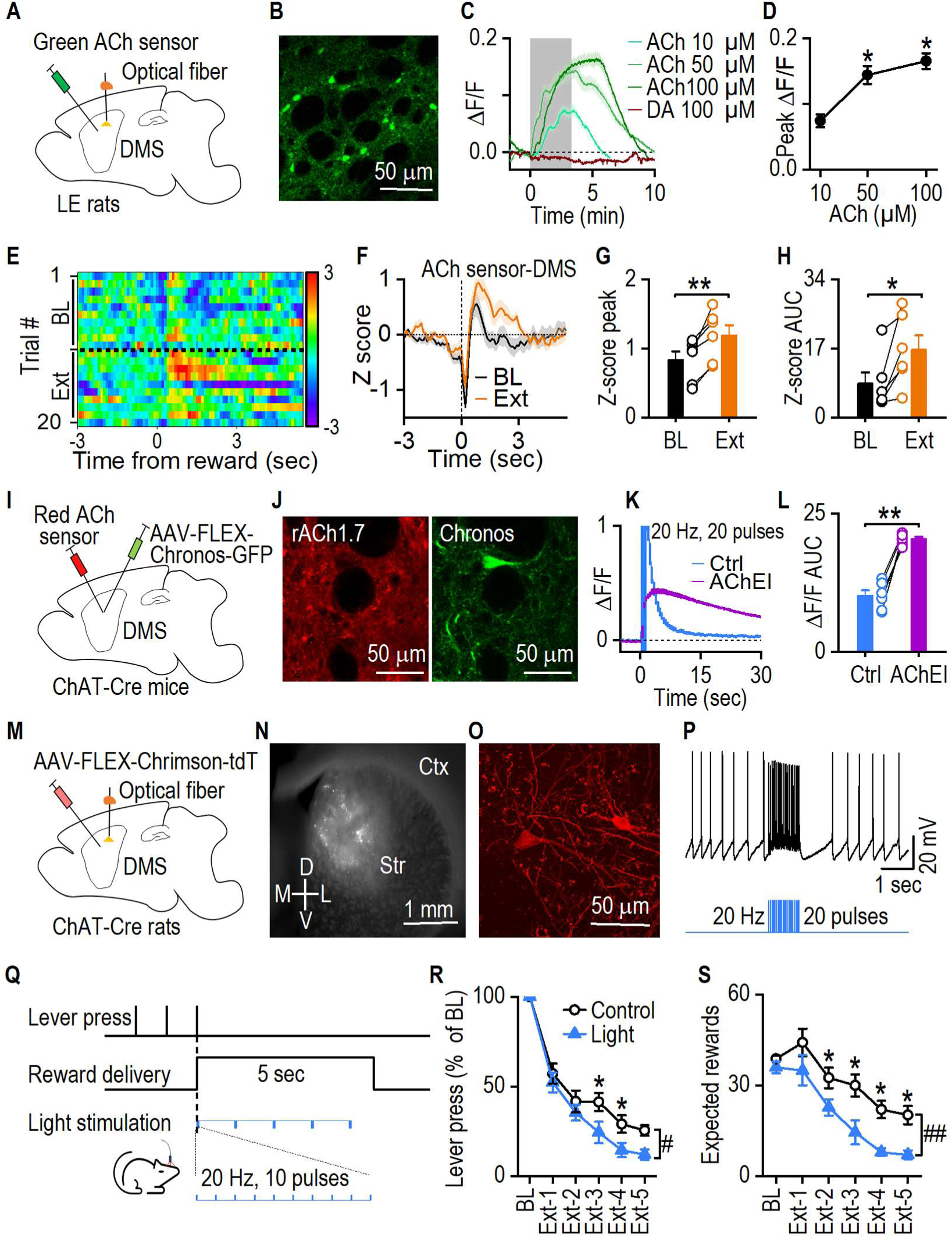
Extinction training enhances ACh release, and augmented CIN burst firing accelerates extinction learning. **A**. Schematic illustrating virus infusion and fiber photometry recording of ACh signal in the DMS. **B**. Sample image of AAV-iAChSnFR expression in the DMS. **C.** Validation of the iAChSnFR sensor using *ex vivo* confocal imaging. Bath application of ACh increased the sensor fluorescence signals, while dopamine had no effect. **D**. Dose response of ACh-induced changes in iAChSnFR signals. \**p* < 0.05 by paired *t* test comparing to 10 µM, n = 6 ROI (region of interest) from two rats. **E**. *In vivo* measurements of ACh release during baseline (BL) and extinction (Ext) training within the same session. Heatmap shows increased ACh signals after reward in extinction trials. **F**. Representative ACh signal traces during BL and Ext. **G, H.** Greater ACh signal peaks and area under curve (AUC) during extinction than BL; \**p* < 0.05, \*\**p* < 0.01 by paired *t* test, n = 6 rats. **I.** Schematic of viral injections. ChAT-Cre mice were injected with AAV-FLEX-Chronos-GFP and AAV-GRAB_rACh1.7_ in the DMS. **J.** Sample images of red ACh sensor expression and GFP fluorescence in the DMS. **K.** Representative traces of ACh sensor fluorescence signals. Optical stimulation of CINs induced a fluorescent signal increase (Ctrl: control). Bath application of AChE inhibitor (AChEI: Tacrine, 100 µM) prolonged the fluorescent decay. **L.** Summary data quantify the area under the curve (AUC); \*\**p* < 0.01 by paired *t* test, n = 6 ROI from two mice. **M.** ChAT-Cre rats received a bilateral infusion of AAV-FLEX-Chrismon-tdT and optical fibers implantation in the DMS. **N.** The expression of AAV-FLEX-Chrimson-tdT in the DMS. **O.** Sample confocal image of Chrimson-tdTomato expressed CINs. **P.** Sample current-clamp recording trace of CIN firing in response to light stimulation. **Q.** The optical stimulation protocol was employed during extinction training. Lever presses triggered both reward delivery and synchronized light stimulation. Light stimulation comprised 5 repetitions of 20 Hz light bursts (590 nm, 10 pulses, 5 ms per pulse) with 0.8 second intervals. Actual rewards were omitted during extinction. **R.** Lever presses of the light-stimulation group were significantly lower than the control group during extinction training. The data were normalized to their baseline lever presses. **S.** The expected rewards were significantly fewer in the light group than in the control group. Two-way RM ANOVA followed by Tukey *post-hoc* test, ^#^*p* < 0.05, ^##^*p* < 0.01; \**p* < 0.05, n = 13 rats (Control) and 13 rats (Light).

CIN burst firing can prompt phasic ACh release. To confirm this, we infused AAV-FLEX-Chronos-GFP and red ACh sensor AAV-GRAB_rACh1.7_ into the DMS of ChAT-Cre mice, leading to selective expression of Chronos in CINs and AAV-GRAB_rACh1.7_ in striatal neurons (Fig. 1I, J). Blue light stimulation (20 Hz, 20 pulses) of CINs increased Ach sensor fluorescence, the duration of which was enhanced by an acetylcholinesterase inhibitor (Fig. 1K, L). Given our observation of heightened ACh release during extinction, we posited that inducing CIN burst firing could facilitate extinction. To verify this, we bilaterally infused AAV-FLEX-Chrimson-tdT into the DMS of ChAT-Cre rats (Fig. 1M). This infusion led to the strong expression of AAV-FLEX-Chrimson-tdT within the DMS (Fig. 1N, O). *Ex vivo* slice recording revealed that yellow light stimulation (20 Hz, 20 pulses) induced burst-pause response in tdTomato-positive CINs (Fig. 1P). After recovering from surgery, rats were trained to press levers for rewards, progressing from fixed ratio (FR) protocols FR1 and FR3 to random ratio (RR) protocols RR5 and RR10 (Supplemental Fig. 1A). Subsequently, they underwent five consecutive daily extinction sessions using RR5. During extinction, the “light” group received reward-timed light stimulation, composed of five light bursts 0.8 seconds apart, each at 20 Hz and with 10 pulses per burst (Fig. 1Q). The control group received no light stimulation. Notably, the light group extinguished faster, with significantly fewer lever presses (Fig. 1R, *F*_(1, 24)_ = 5.19, *p* < 0.05) and expecting fewer undelivered rewards (Fig. 1S, *F*_(1, 24)_ = 15.18, p < 0.01). These changes were not due to general impacts on motor function or motivation by light stimulation, as CIN burst stimulation did not reduce lever presses and earned rewards when rewards were available compared to the control group (Supplemental Fig. 2A, B) or compared to themselves before light stimulation (Supplemental Fig. 2C). Together, these results suggest that CIN burst firing accelerates the extinction of goal-directed learning.

### Thalamically driven CIN excitation enhances extinction learning

We next sought to identify the source of CIN excitation during extinction training. Given that CINs receive substantial thalamic projections from the PfN [29, 36, 37] and the thalamus’ known responsiveness to salient stimuli [30], we postulated the thalamus could be a potential excitatory input source to CINs during extinction. To verify this, we infused AAV-GCaMP6s into the PfN of Long-Evans rats (Fig. 2A). We observed the expression of AAV-GCaMP6s in the PfN (Fig. 2B) and robust responses to electrical stimulations (Fig. 2C). Animals were trained as in Figure 1E, and fiber photometry was used to record thalamic GCaMP6s signals. We discovered that thalamic activity increased around reward omission during extinction, compared to baseline (Fig. 2D, E, F, *t*_(6)_ = -3.31, *p* < 0.05). To further establish the PfN-CIN link, we infused AAV-Chronos-GFP into the PfN and AAV-GRAB_rACh1.7_, into DMS (Fig. 2G). We observed strong expressions of AAV-Chronos-GFP in the PfN (Fig. 2H) and GRAB_rACh1.7_ in the DMS (Fig. 2J). Patch-clamp recordings confirmed that activating thalamic inputs resulted in DMS CIN burst firing (Fig. 2I). Live confocal imaging revealed that this activation also increased DMS ACh release, a signal augmented by an AChE inhibitor and inhibited by glutamatergic antagonists to block PfN-to-CIN transmission (Fig. 2K, L). These findings suggest that thalamic activation can excite CINs and increase ACh release.

**Figure 2.**
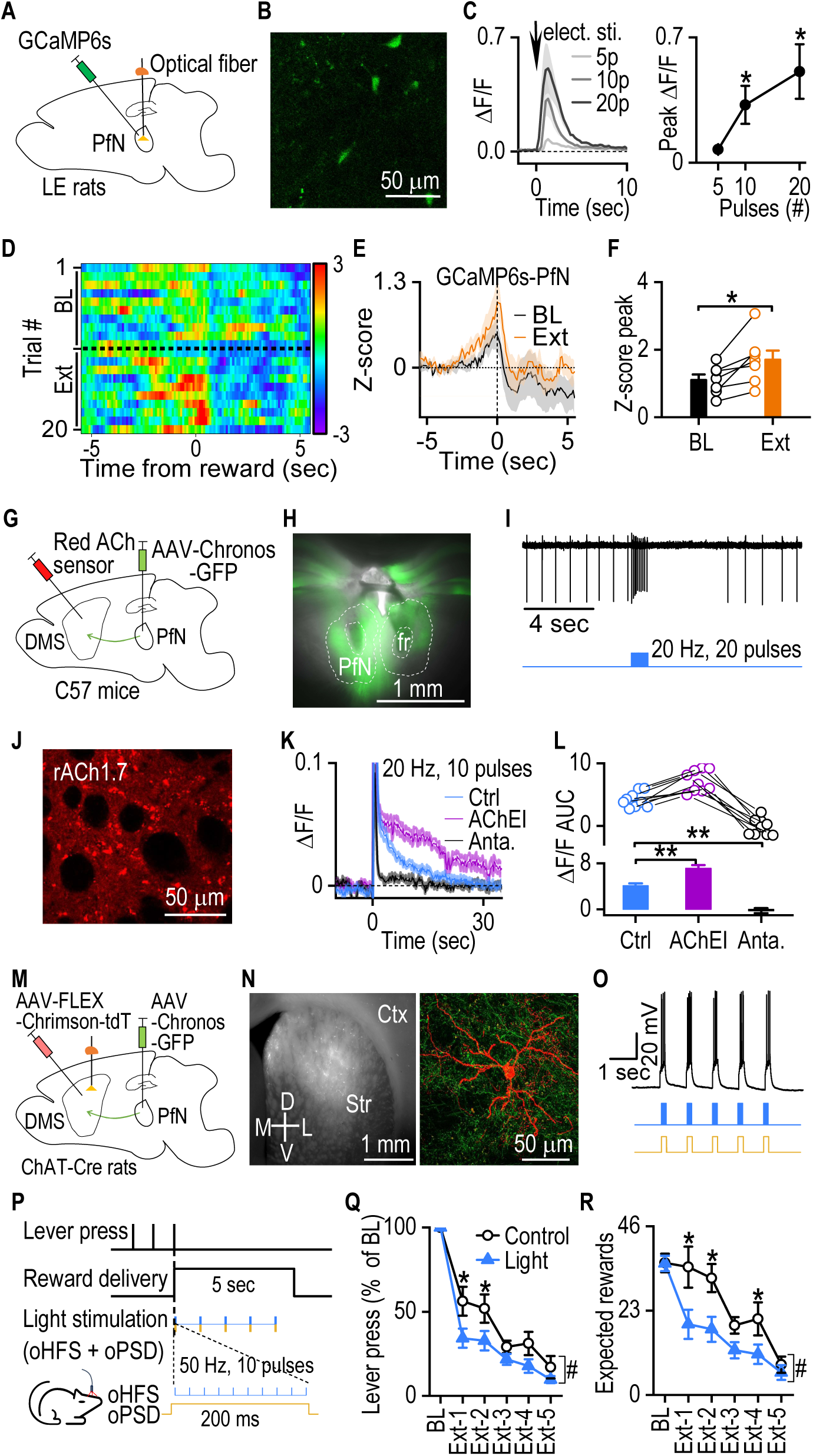
Thalamic-to-CIN pathway activation promotes extinction learning. **A**. Schematic illustrating virus infusion and fiber photometry recording of GCaMP signal in the PfN. **B**. AAV-GCaMP6s expression in the PfN. **C.** Validation of AAV-GCaMP6s using *ex vivo* confocal imaging. Electric stimulation caused an increase in GCaMP signals in PfN slices. \**p* < 0.05 by paired *t* test comparing to 5 pulses, n = 6 ROI (region of interest) from two rats. **D**. *In vivo* measurements of GCaMP signal during baseline (BL) and extinction (Ext) training within the same session. Heatmap shows increased GCaMP signals around reward in extinction trials. **E**. Representative GCaMP signal traces during BL and Ext. **F**. Summary data quantify the peak GCaMP signals; \**p* < 0.05 by paired *t* test, n = 7 rats. **G.** Schematic of viral injections. We infused AAV-GRAB_rACh1.7_ into the DMS and AAV-Chronos-GFP into the PfN of C57BL/6J mice. **H.** Sample image of GFP fluorescence in the PfN. **I.** Optical stimulation of axon terminals from PfN induced a burst firing of CIN in the DMS. **J.** Sample image of rACh1.7 red fluorescence in the DMS. **K.** Representative ACh sensor signal traces. Optical stimulation of axon terminals from PfN induced a fluorescent signal increase in the DMS (Ctrl: control). This increase was diminished with the existence of glutamatergic antagonists (Anta.: NBQX 10 µM + APV 50 µM) and prolonged with the application of AChE inhibitor (AChEI: Tacrine, 100 µM). **L.** Summary data quantify the area under the curve (AUC); \*\**p* < 0.01 by paired *t* test, n = 9 ROI from three mice. **M.** Schematic of viral injection and optical fiber implantation. **N.** Sample image of AAV-FLEX-Chrimson-tdT expression in the DMS and confocal image of Chrimson-tdTomato expressed CIN with Chronos-GFP thalamic inputs. **O.** Sample current-clamp recording trace of CIN firing in response to five repeats of dual light stimulation (470 nm, 50 Hz, 10 pulses; 590 nm, 200 ms). **P.** The optical stimulation protocol was used during the extinction training. Lever presses triggered both reward delivery and synchronized light stimulation. Light stimulation contained five repeats of dual light stimulus within a 5-second reward delivery period. Each repeat consisted of optogenetic high-frequency stimulation (oHFS; 473 nm, 50 Hz, 10 pulses) and optogenetic postsynaptic depolarization (oPSD; 590 nm, 200 ms). Actual rewards were omitted during extinction. **Q.** lever presses during the extinction training. The data were normalized to the percentage of their baseline lever press values. **R.** The expected rewards during the extinction training. Two-way RM ANOVA followed by Tukey *post-hoc* test, ^#^*p* < 0.05, \**p* < 0.05. n = 8 rats (Control) and 9 rats (Light).

If the PfN-CIN pathway is activated during extinction, its heightened activation should promote extinction. To explore this, we infused AAV-Chronos-GFP into the PfN and AAV-FLEX-Chrimson-tdT into the DMS of ChAT-Cre rats, allowing thalamic inputs to express Chronos-GFP and CINs to express Chrimson-tdTomato (Fig. 2N). With *ex vivo* slice recording, the light stimulation protocol designed to induce long-term potentiation (LTP) [37, 38] triggered bursts firing in CINs (Fig. 2O). After recovering from surgery, rats were trained as in Figure 1 (Supplemental Fig. 1B). During extinction, the light group received LTP light stimulation synchronized with reward delivery (Fig. 2P). We used this LTP protocol rather than just presynaptic thalamic stimulation to strengthen PfN-to-CIN transmission selectively [37]. The control group did not receive light stimulation. The light group showed quicker extinction, marked by reduced lever presses (Fig. 2Q, *F*_(1, 15)_ = 7.43, *p* < 0.05) and fewer anticipated rewards (Fig. 2R, *F*_(1, 15)_ = 7.67, *p* < 0.05). These results indicate that PfN-CIN activation promotes the extinction of goal-directed learning.

### Pause in CIN firing is critical for the reversal of goal-directed learning

Burst firing of CINs is typically followed by a pause (Fig. 1L, 2I). We noticed that the ACh signal exhibited an initial drop before a rise and then another dip in the operant self-administration experiment (Fig. 3A). We attribute the ACh signal decrease to the CIN firing pause. However, little is known about the potential behavioral implications of the pause phenomenon. It has been suggested that the pause in CIN firing could provide a time window for corticostriatal plasticity to occur [31, 32], which is pivotal for new learning. Thus, we postulated that a pause is required to update learning. To test this idea, we used optogenetic methods to manipulate the CIN pause, resulting in light-induced pause disruptions (Fig. 3B, C; Supplemental Fig. 3). AAV-FLEX-Chrimson-tdT was infused into the DMS of ChAT-Cre rats (Fig. 3D) for pause disruption. We then used a reversal learning paradigm [37, 39] to uncover behavioral impacts. In this setup, animals initially associate a left lever with sucrose and a right lever with food pellets. Later, these associations are swapped (Fig. 3E, H; Supplemental Fig. 1C, D). After each learning stage, a devaluation test assessed the learning of these associations (Fig. 3F). Light stimulation was applied only during reversal learning and was time-locked to reward delivery (Fig. 3G, H).

**Figure 3.**
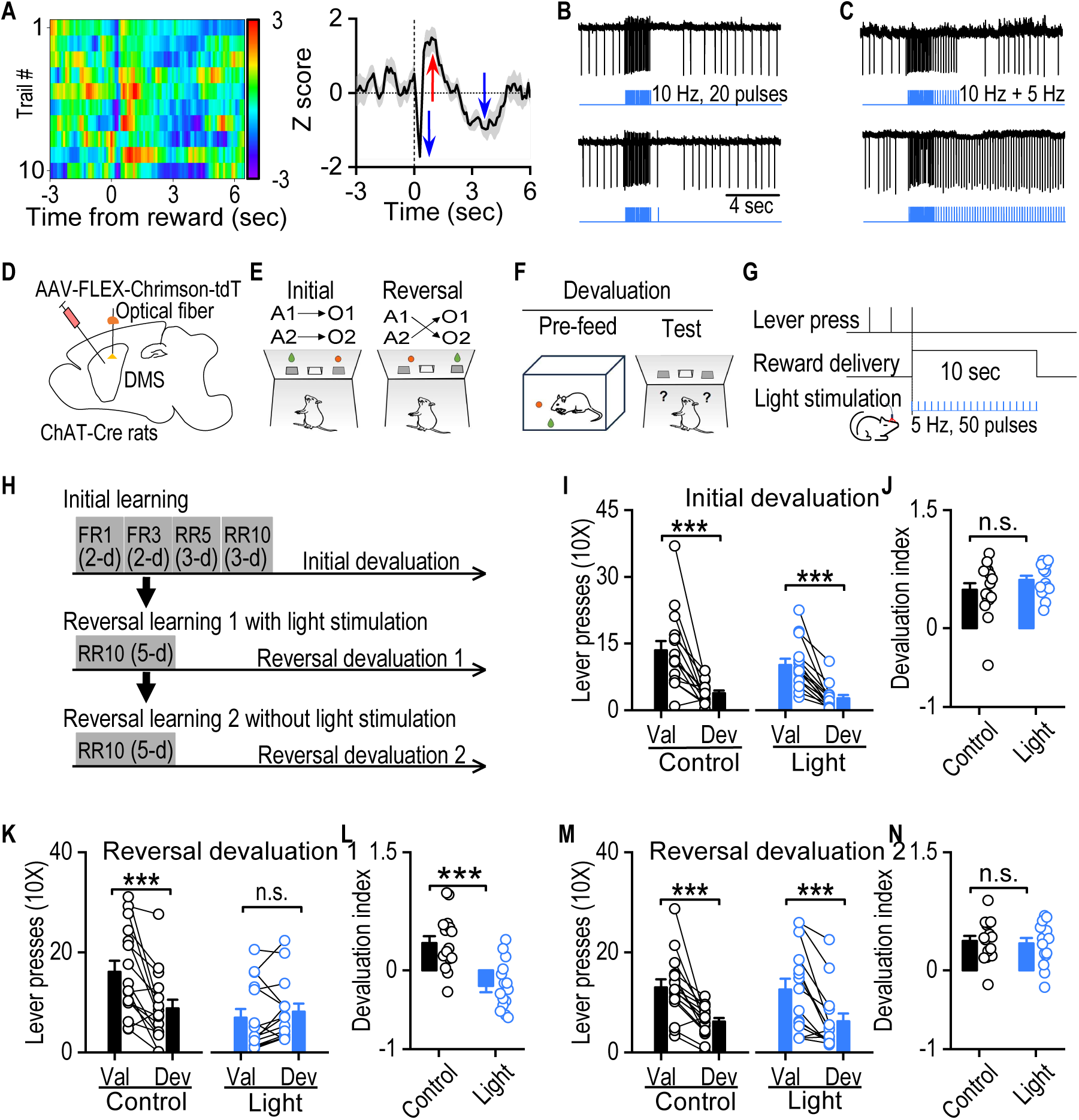
Pause in CIN firing is critical for updating goal-directed learning. **A.** Heat map (left) and trace (right) depict ACh signal variations during training, showing a post-reward decline, rebound, and subsequent drop. **B.** Top: Optical stimulation (10 Hz, 20 pulses) induces burst-pause CIN firing in brain slices of ChAT-Cre;Ai32 mice. Bottom: An action potential can be elicited by an additional optical stimulation during the pause period. **C.** Top: Multiple action potentials can be elicited during the pause period (5 Hz, 10 pulses). Bottom: 5 Hz continuous optical stimulation interrupts CIN firing pause. **D.** Bilateral AAV-FLEX-Chrimson-tdT infusion and optical fiber implantation in the DMS of ChAT-Cre rats. **E.** Rats underwent instrumental learning to press the left lever (A1) for sucrose solution (O1) and the right lever (A2) for food pellets (O2). This training was followed by reversal learning, where A1 led to O2 and A2 led to O1. **F.** Schematic of the devaluation test. Rats were pre-fed with one of the rewards before a 10-minute extinction test. **G.** Optical stimulation protocol used during reversal learning. Rats pressed the lever to receive rewards and synchronized light stimulation (590 nm, 5 Hz, 50 pulses for 10 sec). **H.** Schematic of behavioral training. Rats first underwent initial learning progressed from fixed ratio (FR) protocols FR1, FR3 to random ratio (RR) protocols RR5, RR10 followed by an initial devaluation test. Then, there were two rounds of reversal learning with or without light intervention, each followed by a devaluation test. **I.** Outcome-specific devaluation testing showed both groups pressed the devalued (DeV) lever significantly less than the valued (Val) post-initial learning; \*\*\**p* < 0.001. **J.** Devaluation index, calculated as (Val – DeV) / (Val + DeV), showed no significant difference between groups; *p* = 0.22. **K.** Post-reversal learning devaluation showed decreased DeV lever interaction in the control group, whereas this devaluation effect was not observed in the light group, suggesting impaired reversal learning; \*\*\**p* < 0.001 (Control); *p* = 0.39 (Light). **L.** The devaluation index was significantly lower in the light group than in the control group; \*\*\**p* < 0.001. **M.** Post-reversal learning without light stimulation devaluation test showed that both groups learned reversed action-outcome links, which was evident by more Val lever presses than DeV lever; \*\*\**p* < 0.001. **N.** There is no significant difference in the devaluation index between groups; *p* = 0.73. Mixed model ANOVA was followed by a simple effects test for (I, K, M), an unpaired *t* test for (J, L, N), n = 16 rats (Control) and 15 rats (Light). n.s., not significant.

Upon initial devaluation, both the control and light groups exhibited notably diminished presses on the outcome-satiated devalued lever (Dev_Lever_) (Fig. 3I). There was a main effect of devaluation (*F*_(1,29)_ = 53.68, *p* < 0.001), but no group x devaluation interaction (*F*_(1,29)_ = 0.85, *p* = 0.36). The devaluation index, calculated as (Val_Lever_ - Dev_Lever_) / (Val_Lever_ + Dev_Lever_), showed no significant difference between the groups (Fig. 3J; *t*_(29)_ = -1.25, *p* = 0.22). These results suggested that both groups acquired initial action-outcome contingencies. During reversal devaluation, the control group continued to favor the valued lever (Val_Lever_), whereas the light group indiscriminately pressed on both Val_Lever_ and Dev_Lever_ (Fig. 3K). Statistical analysis revealed a significant group x devaluation interaction (*F*_(1,29)_ = 16.87, *p* < 0.001). Further simple effects test revealed that while the control group had significantly higher valued versus devalued lever presses (Fig. 3K left; *F*_(1,29)_ = 25.01, *p* < 0.001), the light group did not (Fig. 3K right; *F*_(1,29)_ = 0.76, *p* = 0.39). Notably, the light-stimulated rats had significantly lower devaluation indices than controls (Fig. 3L; *t*_(29)_ = 4.66, *p* < 0.001). These results suggest that control rats acquired the reversed action-outcome contingencies, but light-stimulated rats failed to do so. To test the reversible nature of this disruption, all rats underwent further reversal learning without light stimulation (Supplemental Fig. 1D). Following this training phase, reversal devaluation test 2 was performed (Fig. 3H). As shown in Figure 3M, the light group rats recovered their reversal learning abilities, indicating more valued lever pressing than devalued lever pressing. There was a main effect of devaluation (*F*_(1,29)_ = 42.91, *p* < 0.001), but no group x devaluation interaction (*F*_(1,29)_ = 0.07, *p* = 0.80). Their devaluation indices were comparable to controls (Fig. 3N, *t*_(29)_ = 0.35, *p* = 0.73). Further analysis of devaluation indices confirmed the reversibility of light disruption (Supplemental Fig. 4A-C).

Our study observed a notable increase in ACh signal during extinction (Fig. 1E-H), likely stemming from enhanced thalamic activation in CINs (Fig. 2D-L). During the reversal learning, the switches of action-outcome contingencies could also serve as salient stimuli for animals. We predicted that the patterns of ACh signaling and thalamic activity during reversal learning would resemble those observed during extinction. Indeed, when compared to the initial learning session, the reversal session exhibited elevated ACh signal following reward delivery (Supplemental Fig. 5A-D) and amplified thalamic activity (Supplemental Fig. 5E-G). This is consistent with a recent study that showed that reversal learning increased the burst firing of DMS CINs [40].

Taken together, our results suggest that reversal leaning increases PfN-to-CIN transmission and ACh release, followed by a pause in CIN firing, which is required to reverse goal-directed behavior.

### Chronic alcohol consumption reduces burst-pause responses of CINs and impairs reversal of goal-directed learning

Through optogenetic manipulations, we demonstrated that CIN burst firing promotes the extinction of goal-directed learning, while CIN pause is vital for updating it. However, we wanted to investigate if this holds in a more physiological-related condition. Our previous study revealed that chronic alcohol intake and withdrawal led to a long-lasting reduction of thalamostriatal inputs to DMS CINs and impaired reversal learning [37]. Since thalamic inputs trigger burst-pause responses in CINs [27] (Fig. 2I), we postulated that chronic alcohol exposure might modify these CIN responses due to altered thalamostriatal transmission. We explored the effect of long-term alcohol consumption on CINs’ burst-pause responses by using VGluT2-Cre;Ai32;ChAT-eGFP mice (Fig. 4A). Previous studies in VGluT2-Cre mice demonstrated that VGluT2-expressing inputs to the striatum primarily originate from the thalamus [41, 42]. Mice were trained to consume 20% alcohol for 8 weeks using an intermittent-access 2-bottle choice drinking procedure [37, 43]. Striatal slices were prepared either 1 day or 21 days after the last alcohol exposure. We then conducted cell-attached recording of CINs in response to optical stimulation of thalamic inputs (470 nm, 20 Hz, 20 pulses, Fig. 4B). Notably, burst responses to thalamic stimulation gradually decreased as alcohol withdrawal progressed (Fig. 4C, *F*_(2,50)_ = 4.65, *p* < 0.05). Interestingly, a similar trend was observed in the pause responses of CINs, where pause durations shortened over time (Fig. 4D, *F*_(2,50)_ = 5.67, *p* < 0.01). To corroborate the alcohol-associated suppression of thalamostriatal induced burst-pause responses of CINs, AAV-FLEX-Chrimson-tdT was infused into the PfN of thalamus of VgluT2-Cre;ChATeGFP mice (Fig. 4E.). CINs were recorded as above in the cell-attached mode (Fig. 4F), yielding consistent results with above transgenic mice findings (Burst: Fig. 4G, *F*_(2,147)_ = 6.59, *p* < 0.01; Pause: Fig. 4H, *F*_(2,147)_ = 12.45, *p* < 0.001). To provide further validation, CINs were also recorded in whole-cell current clamp mode (Fig. 4I), confirming the progressive reduction of burst-pause responses with ongoing alcohol withdrawal (Burst: Fig. 4J, *F*_(2,16)_ = 6.25, *p* < 0.01; Pause: Fig. 4K, *F*_(2,16)_ = 8.29, *p* < 0.01).

**Figure 4.**
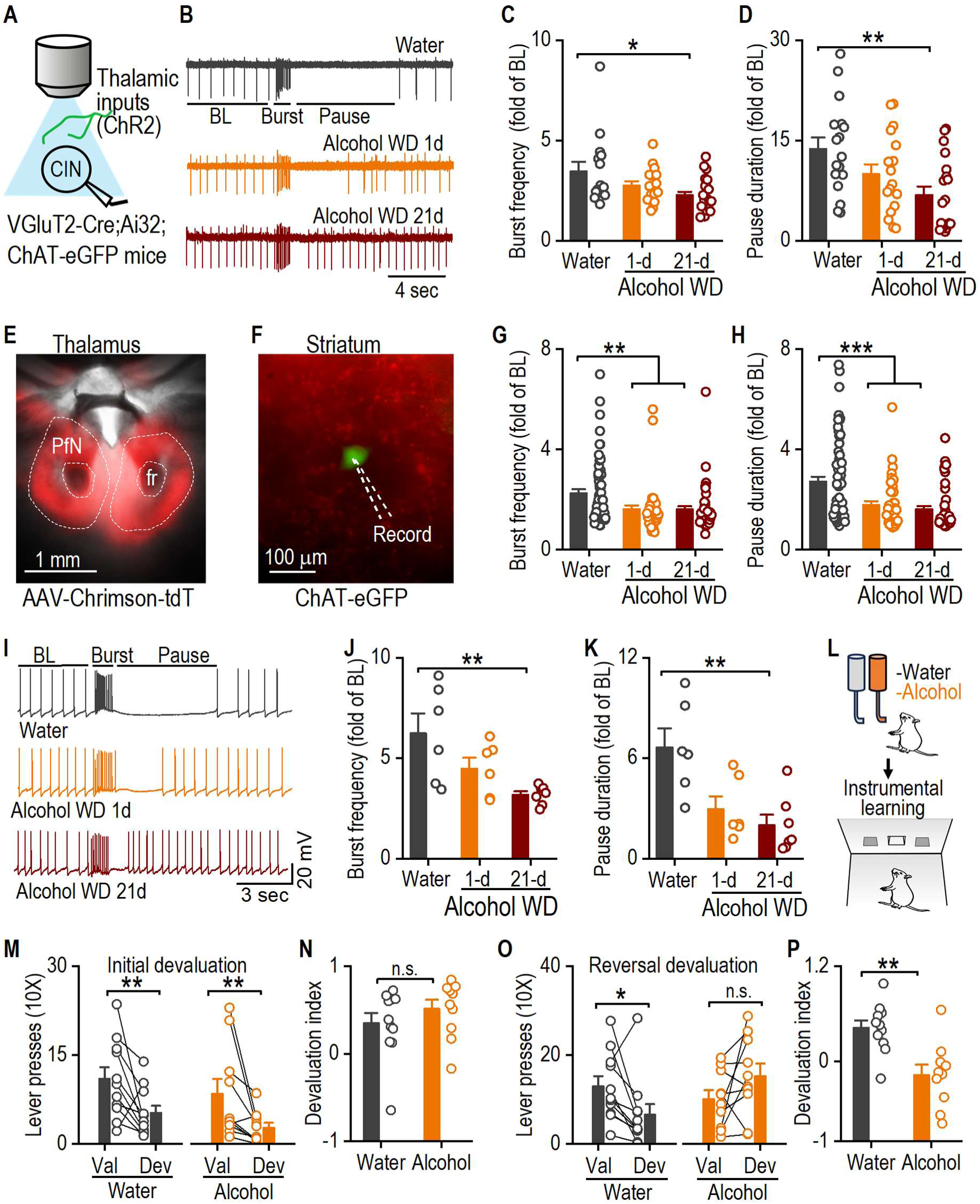
Excessive alcohol intake causes a long-lasting reduction of thalamically evoked burst-pause response in DMS CINs and impairs reversal of instrumental learning. **A**. Schematic of light stimulation of ChR2-expressing thalamic inputs and whole-cell recording of CINs in VGluT2-Cre;Ai32;ChT-eGFP mice. **B**. Sample traces from cell-attached recording of CINs in response to optical stimulation of thalamic inputs (470 nm, 20 Hz, 20 pulses). **C**. Bar graph of burst frequency expressed as fold changes of baseline firing frequency. \**p* < 0.05. **D**. Bar graph of pause duration expressed as fold changes of baseline interspike intervals. \*\**p* < 0.01. **E.** Sample image showing tdTomato fluorescence in the PfN. fr, fasciculus retroflexus. **F.** Sample image of tdTomato fluorescence and ChAT-eGFP neuron in the striatum. **G**. Same as panel C. \*\**p* < 0.01. **H**. Same as panel D. \*\*\**p* < 0.001. **I.** Sample traces from whole-cell recording of CINs in response to optical stimulation of thalamic inputs (470 nm, 20 Hz, 20 pulses). **J**. Same as panel C. \*\**p* < 0.01. **K**. Same as panel D. \*\**p* < 0.01. **L**. Schematic of treatment and behavioral training. The Long-Evans rats were introduced to consume 20% alcohol for 8 weeks using an intermittent-access 2-bottle choice drinking procedure and then trained for instrumental learning, as in Figure 3E. **M.** Both water and alcohol groups pressed the devalued (DeV) lever significantly less than the valued (Val) lever after initial learning; \*\**p* < 0.01. **N.** Devaluation index, (Val - DeV) / (Val + DeV), revealed no significant difference between groups; *p* = 0.31. **O.** Post-reversal learning devaluation showed decreased DeV lever interaction in the water group, whereas this devaluation effect was not observed in the alcohol group, indicating impaired reversal learning; \**p* < 0.05 (Water); *p* = 0.09 (Alcohol). **P.** Alcohol group had a significantly lower devaluation index than the water group; \*\**p* < 0.01. One-way ANOVA followed by Tukey *post-hoc* test for (C, D): n = 15 neurons from 5 mice (Water 15/5), (Alcohol WD 1-d 16/5), (Alcohol WD 21-d 22/5); for (G,H): (Water 65/10), (Alcohol WD 1-d 41/7), (Alcohol WD 21-d 44/7); for (J, K): (Water 6/3), (Alcohol WD 1-d 6/3), (Alcohol WD 21-d 7/3). Mixed model ANOVA was followed by a simple effects test for (M,O), an unpaired *t* test for (N, P), n = 11 rats (Water) and 10 rats (Alcohol). BL, baseline; WD, withdrawal; n.s., not significant.

To test the behavioral consequences of these reduced burst-pause responses of CINs, Long-Evans rats were exposed to water (control) or 20% alcohol for 8 weeks (Fig. 4L). The animals were then trained for instrumental learning, as in Figure 3, including both initial (Supplemental Fig. 1E) and reversal learning (Supplemental Fig. 1F). The initial devaluation test showed that both the water and alcohol groups acquired the initial action-outcome contingencies (Fig. 4M). Statistical analysis showed a main effect of devaluation (*F*_(1,19)_ = 14.27, *p* < 0.01), but no group x devaluation interaction (*F*_(1,19)_ < 0.001, *p* = 0.99). Analysis of the devaluation index identified no significant difference between groups (Fig. 4N, *t*_(19)_ = -1.04, *p* = 0.31). However, after reversal learning, the alcohol group preferred the devalued lever (previously the valued lever) (Fig. 4O), indicating the initial action-outcome contingencies were still present. Statistical analysis revealed a significant group x devaluation interaction (*F*_(1,19)_ = 8.57, *p* < 0.01). Further simple effects test revealed that while the water group had significantly higher valued versus devalued lever presses (Fig. 4O left; *F*_(1,19)_ = 5.53, *p* < 0.05), the alcohol group did not (Fig. 4O right; *F*_(1,19)_ = 3.26, *p* = 0.09). Furthermore, the devaluation index of the alcohol group was significantly lower than the control group (Fig. 4P; *t*_(19)_ = 3.79, *p* < 0.01). These results indicated that while water controls acquired new action-outcome contingencies after reversal learning, alcohol-exposed rats failed to update the reversed contingencies.

## Discussion

In this study, we observed elevated ACh release in the DMS during both extinction and reversal learning. We also found increased thalamic activities during these learning phases. The changes of action-outcome contingencies during extinction and reversal learning could serve as salient stimuli to stimulate thalamic neuronal activity, which, in turn, activates CINs, leading to increased ACh release. With optogenetic manipulations, we showed that burst firing of CIN promoted extinction and that the pause in CIN firing was required for reversal learning. These findings demonstrate the significance of CIN burst firing and pause response in the extinction and updating goal-directed learning.

We demonstrated that burst firing of CIN promotes extinction. Research has indicated that CIN activity can influence the extinction of cocaine contextual memories [44]. Our findings suggest that the burst firing of CINs facilitates the extinction of goal-directed learning. This modulation is likely through the neurotransmitter ACh influencing downstream targets. The striatum has the highest activity of acetylcholinesterase (AChE, the enzyme degrading acetylcholine) in the brain [13, 16, 45], indicating tight control over ACh levels. As the primary ACh source in the striatum [16], CIN burst firing can elevate striatal ACh levels. ACh can have distinct modulations on different striatal cell types through different receptors. For example, via M4 muscarinic receptors (M4Rs), it can suppress excitatory inputs to dopamine D1 receptor-expressing medium spiny neurons (D1-MSNs) [37], promoting long-term depression (LTD) in these neurons [46]. Notably, M4Rs are known to regulate D1-MSN function negatively [47]. Through M1 muscarinic receptors (M1Rs), ACh can heighten short-term excitability in dopamine D2 receptor-expressing MSNs (D2-MSNs) [27, 37]. Considering the striatal D1-MSNs and D2-MSNs lead to the “Go” and “No-Go” pathways, respectively, CIN activation might aid extinction by suppressing the “Go” pathway and enhancing the “No-Go” pathway. This aligns with research showing enhanced D2-MSN activation during extinction training [48, 49]. Another mechanism through which ACh could promote extinction involves activating GABAergic interneurons in the striatum through nicotinic acetylcholine receptor (nAchR) [50, 51]. The activation of these GABAergic interneurons could exert robust modulation over striatal MSNs’ activity and control the functioning of striatal circuities [52, 53]. However, while CIN activity can modulate the extinction process, it may not be required for extinction. Inhibiting CIN activity resulted in only a slight delay of extinction in the first session (Supplemental Fig. 6), and our previous study showed a similar effect. When CINs were inhibited, rats pressed more levers only in the first 10-20 minutes of the first 40-minute extinction session [39]. These results imply that CINs predominantly modulate extinction rather than being required for it. Dopamine is most likely the primary signal for extinction, serving as a key modulator of corticostriatal synaptic plasticity [54]. This is supported by previous findings, which show that inhibiting dopamine neurons mimics negative reward prediction error and promotes the extinction of learned associations [55–58].

Pause in CIN firing is critical for the reversal of goal-directed learning. CIN tonic firing maintains a consistent ACh baseline level in the striatum. This ACh generally serves as an inhibitory signal, suppressing corticostriatal transmission via presynaptic M2 receptors [27] and inhibiting MSNs through GABAergic interneurons [50–53]. When CINs pause their firing, ACh release stops and is quickly cleared due to the rapid catabolic activity of AChE [59]. This ACh-free period, combined with dopamine signaling [17, 22, 23, 31], is crucial for striatal-dependent learning. A recent study showed that for long-term corticostriatal plasticity to occur, dopamine increase, CIN pause, and MSN depolarization must synchronize [32]. This suggests that the pause in CINs firing is critical for neuronal plasticity to occur and, hence, could play an essential functional role in changing animal behavior. Disrupting this pause showed that it’s vital for updating goal-directed learning. Animals with chronic alcohol exposure had shorter CIN pauses, leading to impaired reversal learning. Notably, initial learning wasn’t affected. This suggests the pause duration may also have essential implications on behavioral functions. Indeed, prior studies indicate varying CIN pause durations based on stimuli type, with shorter pauses responding to aversive stimuli than to appetitive one [60]. The impaired reversal learning in alcohol-exposed animals may stem from both reduced burst and pause responses. The former hinders old learning extinction, while the latter affects new learning update. Indeed, during the devaluation test after reversal learning alcohol group overall still pressed more devalued lever (which is the previous valued lever) than the valued lever, indicating that the previous action-outcome contingencies had not been fully extinguished. This suggests that CIN burst-pause dynamics may work cooperatively under normal conditions: burst firing aids in old learning extinction, while the subsequent pause assists with new learning. These mechanisms underscore behavioral adaptability. As for alcohol’s impact on CIN responses, our previous study indicated it’s unlikely due to changes in presynaptic release because of the unaffected paired-pulse ratio (PPR) [37]. We suspect postsynaptic changes play a role, evidenced by decreased AMPA-induced currents in alcohol-exposed CINs (Supplemental Fig. 7A, B), indicating a change in postsynaptic AMPA receptor functions. Another possible factor affecting thalamically induced burst-pause responses is CIN’s baseline firing frequency. Here, we replicated our finding [37] that chronic alcohol intake and withdrawal increased the spontaneous firing of CINs (Supplemental Fig. 7E). Optimal ACh signaling is vital for attention and learning [61], and there appears to be an ideal CIN activity level in the striatum for modulating dynamic responses. Any deviation from this baseline may compromise their adaptability to salient behavioral cues.

CINs display very dynamic responses during behaviors [25, 26]. During our instrumental learning paradigm, we noted CINs exhibiting multiphasic responses. Typically, during reward delivery, the ACh signal dipped, surged, and then dipped again. What causes these changes in ACh signals is not clear. Based on previous literature, we believe the initial dip might result from GABA-mediated CIN inhibition [25, 26], as CINs receive multiple inhibitory GABAergic inputs, including those from striatal MSNs [62–64] and can self-inhibit by activation of GABAergic interneurons through nAChR [50, 52]. Here, no preceding CIN excitation was seen for this dip. We hypothesize that it is triggered by GABA release from D1-MSNs during reward-acquiring actions, with supporting evidence from our observation that D1-MSN optogenetic activation can induce a pause-rebound in CIN firing [39]. The subsequent ACh increase may have dual origins: pause-rebound firing of CINs [25, 39] and thalamic CIN activation [27, 30]. The following ACh dip might arise from another CIN firing pause, often seen in post-thalamic activation [27, 65]. Self-initiated recurrent inhibition of CINs [50, 52] could also play a role here. Moreover, this pause response could be modulated by dopamine signals [27, 66–68]. Although the mechanisms for the multiphasic responses of CINs are not entirely clear, this pause and rebound, followed by a second pause of CINs, has been repeatedly observed in behavioral animals [60, 69, 70]. Future studies are needed to elucidate the mechanisms of CIN dynamic responses, dissect out which pause response is critical for updating goal-directed learning, and explore how various signals collaborate during the pause to enhance neural plasticity in behavior.

In summary, our study discovered that thalamic-driven CIN burst firing promotes extinction learning, while the pause in CIN firing is pivotal for reversal learning of goal-directed behavior. A deeper understanding of the behavioral implications of CIN dynamic responses would pave the way to elucidate their critical functions in both the healthy and diseased brain.

## Methods

### Animals

ChAT-eGFP (stock # 007902), VGluT2-Cre (stock # 016963), and Ai32 (stock # 012569) were purchased from the Jackson Laboratory. All mice were backcrossed onto a C57BL/6J background. VGluT2-Cre mice were crossed with Ai32 to generate VGluT2-Cre;Ai32 line. VGluT2-Cre;Ai32 mice were crossed with ChAT-eGFP to generate VGluT2-Cre;Ai32;ChAT-eGFP triple transgenic mice. Both male and female mice were used for electrophysiology studies. Male Long-Evans rats (3 months old) purchased from Harlan Laboratories were used for behavioral testing. Long Evans-Tg(ChAT-Cre) rats were purchased from Rat Resource & Research Center (stock# 00658). ChAT-Cre rats were bred in-house. Both male and female rats (3 months old) were used for behavioral testing. Animals were housed individually at 23°C under a 12-h light:dark cycle, with lights on at 7:00 A.M. Food and water were provided ad *libitum*.

### Reagents

D-APV (Category # ab120003) was purchased from Abcam. DNQX (6,7-dinitroquinoxaline-2,3-dione) (Category # 0189) and Cholinesterase inhibitor (AChEI) (Tacrine hydrochloride, Category # 0965) were purchased from Tocris. All other chemicals for electrophysiological recording were obtained from Sigma. rAAV8/Syn-Chrimson-tdT (#AV5841D), rAAV8/Syn-FLEX-Chrimson-tdT (#AV5844D), and rAAV8/Syn-Chronos-GFP (#AV5842B) were purchased from the University of North Carolina Vector Core. pAAV.CAG.iAChSnFR (#137955-AAV9) and pAAV.Syn.GCaMP6s.WPRE.SV40 (#100843-AAV5) was purchased from Addgene. rAAV/hSyn-GRAB_rACh1.7_ (# PT-5488) was purchased from BrainVTA.

### Behavioral Procedures

#### Intermittent access to 20% alcohol 2-bottle choice drinking procedure

This procedure was conducted as described previously [43, 71–77]. Briefly, animals were given concurrent access to one bottle of alcohol (20%, in water) and one bottle of water for 24-h periods, which were separated by 24- or 48-h periods of alcohol deprivation. Alcohol intake (g/kg/day) was calculated by determining the weight of 20% alcohol solution consumed and multiplying this by 0.2. Water control animals only have access to water.

### Operant Conditioning training

Operant conditioning was conducted on rats as previously described [37, 39]. Blinding was applied to behavioral experiments. An independent observer coded and randomized animals using a computer-generated blinding algorithm. Researchers in the lab trained rats without knowing the treatment plan for the animals. Food was restricted to maintaining 80% of the original body weight of the animals for the duration of behavioral studies.

#### Magazine training

This procedure was adapted from previous studies [7, 37]. After 5 days of food restriction, rats were trained for magazine entries for two consecutive days. During these training sessions, a reinforcer (either a food pellet or 0.1 mL sucrose solution) was delivered along with illumination of the magazine light for 1 sec with a random interval between each reinforcer (on average 60 sec). For each day, rats received either 20 food pellets or 20 sucrose deliveries during the first training session and were then switched to receive the other reward in the second training session. The house light was illuminated throughout the session, and no levers were available during magazine training.

#### Acquisition of initial contingencies

Following magazine training, rats were trained to access different reinforcers via lever pressing over sessions. Each session consisted of 4 blocks (2 blocks per lever), separated by a 2.5-min timeout during which no levers were available, and all lights were extinguished. Only one lever was available during each block (pseudorandom presentation), which lasted for 10 min or until 10 reinforcers had been earned. For extinction sessions, only 2 blocks (one block per lever, 5 min for each block), separated by a 1.5-min timeout, were used. For half of the animals in each group, the left lever was associated with food pellet delivery and the right lever with sucrose solution delivery. The remaining animals were trained using the opposite pairs of action-outcome contingencies. Lever training started with a fixed ratio 1 (FR1) schedule in which every lever press resulted in the delivery of a reinforcer. Some animals may need more training sessions to form the association between lever pressing and reward delivery. After 2 days of stable FR1 training, the training schedule was elevated to FR3 for 2 days. Then, we proceeded to a random ratio 5 (RR5) schedule for the next 3 days, during which a reinforcer was delivered after lever pressing with a probability of 0.2. An RR10 (or a probability of 0.1) training schedule was then employed for 3 days.

#### Devaluation test

After the final RR10 training, devaluation testing was performed for 2 days. On both days, rats were habituated in a dark, quiet room (different from the operant room) for 30 min, then were given *ad libitum* access to either the food pellets (25 g placed in a bowl) or the sucrose solution (100 mL in a drinking bottle) in a devaluation cage for 1 hour. The devaluation cage was similar to their home cage but with new bedding. The rats were then placed in the operant chamber for a 10-min extinction choice test. Both levers were extended during this test, but no outcomes were delivered in response to any lever press. On the second devaluation day, the rats were pre-fed, as described, with the other reward before repeating the same extinction test. If rats fail to perform during the devaluation test, then the pre-feed reward amount and duration need to be tailored to individual animals. Lever presses were recorded, and those on the lever that the rat had learned to associate with the non-devalued reward were termed Val_Lever_, while those on the lever associated with the devalued reward were termed Dev_Lever_. The devaluation index [(Val_Lever_ - Dev_Lever_)/(Val_Lever_ + Dev_Lever_)] was then used to determine the extent of goal-directed versus habitual behavior.

#### Contingency reversal and devaluation testing

After the devaluation test, rats were retrained on their current action-outcome contingencies for 1 day. The contingencies were then reversed so that the lever that previously delivered food now delivered sucrose, and the rats were trained using the RR10 schedule. All other procedures were unchanged. The contingency reversal training lasted for 5 days. The rats then underwent devaluation testing again using the procedure described above.

### Electrophysiology

Slice electrophysiology was performed as previously described [72, 73, 78, 79]. Animals were weaned around postnatal 21 d and consumed 20% alcohol for 6-8 weeks in the intermittent-access 2-bottle choice drinking procedure. Animals were sacrificed 24 h or 21 d after their last alcohol consumption, and 250-µm coronal sections containing the striatum were prepared in an ice-cold cutting solution containing (in mM): 40 NaCl, 148.5 sucrose, 4 KCl, 1.25 NaH_2_PO_4_, 25 NaHCO_3_, 0.5 CaCl_2_, 7 MgCl_2_, 10 glucose, 1 sodium ascorbate, 3 sodium pyruvate, and 3 myoinositol, saturated with 95% O_2_ and 5% CO_2_. Slices were then incubated in a 1:1 mixture of cutting solution and external solution at 32°C for 45 min. The external solution contained the following (in mM): 125 NaCl, 4.5 KCl, 2.5 CaCl_2_, 1.3 MgCl_2_, 1.25 NaH_2_PO_4_, 25 NaHCO_3_, 15 sucrose, and 15 glucose, saturated with 95% O_2_ and 5% CO_2_. Slices were then maintained in an external solution at room temperature until use.

Slices were perfused with the external solution at a flow rate of 3-4 mL/min at 32°C. The CINs in the DMS were identified either by differential interference contrast or by fluorescence. Whole-cell patch-clamp and cell-attached recordings were made using a MultiClamp 700B amplifier controlled by pClamp 10.4 software (Molecular Devices). Optogenetically-evoked CIN firing was induced by light stimulation through the objective lens. Chronos was activated by a 470-nm laser; Chrimson was activated by a 590-nm laser. For cell-attached and whole-cell current-clamp recordings, we used a K^+^-based intracellular solution containing (in mM): 123 potassium gluconate, 10 HEPES, 0.2 EGTA, 8 NaCl, 2 MgATP, 0.3 NaGTP (pH 7.3), with an osmolarity of 270–280 mOsm. Data was analyzed using Clampfit (in pClamp 10.7, Molecular Devices).

#### *In vivo* fiber photometry recording

Fiber photometry was conducted as previously described [39, 80]. Rats were connected to fiber before data collection to acclimate to the fiber patch cable. During the data collection session, the spectrum data was recorded continuously at 10 Hz using OceanView 1.6.7. At the same time, behavior data was collected. 488 nm laser was delivered to excite iAChSnFR or GCaMP6s. The percentage Δ*F/F* was calculated by 100 × (*F* − *F*_mean_)/*F*_mean_, where *F*_mean_ was the mean fluorescence intensity. The Z-score (Z) was calculated using MATLAB by

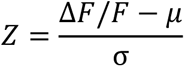

To analyze the fiber photometry data in the context of rat behavior, MATLAB scripts were developed. The code used for the analysis of fiber-photometry data is available online at the Zenodo public repository [https://doi.org/10.5281/zenodo.7948766].

To normalize signals across animals and sessions, we calculated a single-baseline fluorescence value for each trial using the average of the first 5-second period during 10-s before the reward delivery and subtracted that from the signal. Peak amplitudes were calculated by taking the maximum value between 1-s before and 2-s after the reward delivery. The area under the curve (AUC) values were restricted to 0-s to 5-s after the reward delivery time window.

### Stereotaxic surgery and Histology

Stereotaxic viral infusions were performed as described previously [43, 73, 79, 81]. Briefly, animals were anesthetized using isoflurane and mounted in a rodent stereotaxic frame (Kopf). The skin was opened to uncover the skull and expose Bregma and Lambda and the location of the desired injection site. A three-axis micromanipulator was used to measure the spatial coordinates for Bregma and Lambda. Small drill holes were made in the skull at the appropriate coordinates, according to the Paxinos atlas [82, 83]. Two microinjectors were loaded with the virus and then lowered into the appropriate coordinates. The virus was infused into the brain at a rate of 0.1 µL/min. To avoid backflow of the virus, microinjectors were left in place for 10 min after the infusion was complete. Following their removal, the skin was sutured and the animals were allowed to recover for at least 1 week prior to the next experiment. For fiber implantation, following virus injections, bilateral optical fiber implants (300-μm core fiber secured to a 1.25-cm ceramic ferrule with 5 mm of fiber extending past the end of the ferrule) were lowered into 0.1mm above the virus injection sites. Implants were secured on the skull using metal screws and dental cement (Henry Schein) and covered with denture acrylic (Lang Dental). The incision was closed around the head cap, and the skin was vet-bonded to the head cap. The coordinates for mice – DMS (AP, +0.02; ML, ± 1.8; DV, -2.7 mm) for AAV-GRAB_rACh1.7_ sensor (0.4 µL); PfN (AP, -2.2; ML, ± 0.7; DV, -3.5 mm) for AAV-Chronos-GFP (0.5 µL), AAV-Chrimson-tdT (0.5 µL). The coordinates for rats – DMS (AP, 0.0; ML, ± 2.8; DV, -4.85 mm) for AAV-FLEX-Chrimson-tdT(1 µL), AAV-iAChSnFR (1 µL); PfN (AP, -4.2; ML, ± 1.25; DV, -6.2 mm) for AAV-GCaMP6s (1 µL).

The histology procedure was performed as described previously [73, 79, 84]. Briefly, mice were anesthetized and perfused intracardially with 4% paraformaldehyde in phosphate-buffered saline (PBS). Whole brains were taken out and placed into 4% paraformaldehyde in PBS for post-fixation overnight (4°C), then moved to 30% sucrose in PBS (4°C) and allowed to sink to the bottom of the container before preparing for sectioning. Frozen brains were cut into 50-μm coronal sections on a cryostat. A confocal laser-scanning microscope (Fluorview-1200, Olympus) was used to image these sections with a 470-nm laser (to excite eYFP and GFP) and a 593-nm laser (to excite tdT). All images were processed using Imaris 8.3.1 (Bitplane, Zurich, Switzerland).

### Statistical analysis

All data are expressed as mean ± SEM. Statistical analyses were performed using SigmaPlot 12.0 (Systat Software Inc.). Normal distribution was tested; unpaired *t* test, paired *t* test, and one-way ANOVA or two-way RM ANOVA followed by Tukey *post-hoc* test were used to determine statistical significance as appropriate, with an alpha value of 0.05. Mixed model ANOVA followed by a simple effects test was conducted for Fig. 3I, K, M, and Fig. 5M, O in SPSS 29.0 [12].

## Acknowledgments

We appreciate Ms. Valerie Vierkant’s critical comments on our manuscript. This research was supported by NIAAA U01AA025932 (J.W.), R01AA027768 (J.W.), and R01AA030293 (J.W.).

## Author contributions

J.W. conceived, designed, and supervised all the experiments in the study. Z.H. designed and performed electrophysiology experiments and analyzed the data. Z.H. and R.C. designed and performed fiber photometry experiments and analyzed the data. Z.H., X.X., and M.H. designed and performed the behavior experiments and analyzed the data. Z.H. and X.W. conducted histology experiments.

## References

1. Uddin, L.Q., Cognitive and behavioural flexibility: neural mechanisms and clinical considerations. Nat Rev Neurosci, 2021. 22(3): p. 167–179.

2. Stephenson-Jones, M., et al., A basal ganglia circuit for evaluating action outcomes. Nature, 2016. 539(7628): p. 289–293.

3. Stephenson-Jones, M., et al., Independent circuits in the basal ganglia for the evaluation and selection of actions. Proc Natl Acad Sci U S A, 2013. 110(38): p. E3670–9.

4. Kreitzer, A.C. and R.C. Malenka, Striatal plasticity and basal ganglia circuit function. Neuron, 2008. 60(4): p. 543–54.

5. Park, J., L.T. Coddington, and J.T. Dudman, Basal Ganglia Circuits for Action Specification. Annu Rev Neurosci, 2020. 43: p. 485–507.

6. Sharpe, M.J., et al., An Integrated Model of Action Selection: Distinct Modes of Cortical Control of Striatal Decision Making. Annu Rev Psychol, 2019. 70: p. 53–76.

7. Bradfield, L.A. and B.W. Balleine, Thalamic control of dorsomedial striatum regulates internal state to guide goal-directed action selection. J Neurosci, 2017. 37(13): p. 3721–3733.

8. Ragozzino, M.E. and D. Choi, Dynamic changes in acetylcholine output in the medial striatum during place reversal learning. Learn Mem, 2004. 11(1): p. 70–7.

9. Ragozzino, M.E., et al., Acetylcholine activity in selective striatal regions supports behavioral flexibility. Neurobiol Learn Mem, 2009. 91(1): p. 13–22.

10. Matamales, M., et al., Aging-related dysfunction of striatal cholinergic interneurons produces conflict in action selection. Neuron, 2016. 90(2): p. 362–73.

11. Aoki, S., et al., Role of striatal cholinergic interneurons in set-shifting in the rat. J Neurosci, 2015. 35(25): p. 9424–31.

12. Bradfield, L.A., et al., The thalamostriatal pathway and cholinergic control of goal-directed action: interlacing new with existing learning in the striatum. Neuron, 2013. 79(1): p. 153–66.

13. Bolam, J.P., B.H. Wainer, and A.D. Smith, Characterization of cholinergic neurons in the rat neostriatum. A combination of choline acetyltransferase immunocytochemistry, Golgi-impregnation and electron microscopy. Neuroscience, 1984. 12(3): p. 711–8.

14. Witten, I.B., et al., Cholinergic interneurons control local circuit activity and cocaine conditioning. Science, 2010. 330(6011): p. 1677–81.

15. Aosaki, T., et al., Responses of tonically active neurons in the primate’s striatum undergo systematic changes during behavioral sensorimotor conditioning. J Neurosci, 1994. 14(6): p. 3969–84.

16. Woolf, N.J. and L.L. Butcher, Cholinergic neurons in the caudate-putamen complex proper are intrinsically organized: a combined Evans blue and acetylcholinesterase analysis. Brain Res Bull, 1981. 7(5): p. 487–507.

17. Aosaki, T., et al., Acetylcholine-dopamine balance hypothesis in the striatum: an update. Geriatr Gerontol Int, 2010. 10 Suppl 1: p. S148–57.

18. Kimura, M., J. Rajkowski, and E. Evarts, Tonically discharging putamen neurons exhibit set-dependent responses. Proc Natl Acad Sci U S A, 1984. 81(15): p. 4998–5001.

19. Apicella, P., E. Scarnati, and W. Schultz, Tonically discharging neurons of monkey striatum respond to preparatory and rewarding stimuli. Exp Brain Res, 1991. 84(3): p. 672–5.

20. Doig, N.M., et al., Cortical and thalamic excitation mediate the multiphasic responses of striatal cholinergic interneurons to motivationally salient stimuli. J Neurosci, 2014. 34(8): p. 3101–17.

21. Joshua, M., et al., Midbrain dopaminergic neurons and striatal cholinergic interneurons encode the difference between reward and aversive events at different epochs of probabilistic classical conditioning trials. J Neurosci, 2008. 28(45): p. 11673–84.

22. Morris, G., et al., Coincident but distinct messages of midbrain dopamine and striatal tonically active neurons. Neuron, 2004. 43(1): p. 133–43.

23. Cragg, S.J., Meaningful silences: how dopamine listens to the ACh pause. Trends Neurosci, 2006. 29(3): p. 125–31.

24. Chantranupong, L., et al., Dopamine and glutamate regulate striatal acetylcholine in decision-making. Nature, 2023.

25. Schulz, J.M. and J.N. Reynolds, Pause and rebound: sensory control of cholinergic signaling in the striatum. Trends Neurosci, 2013. 36(1): p. 41–50.

26. Zhang, Y.F. and S.J. Cragg, Pauses in Striatal Cholinergic Interneurons: What is Revealed by Their Common Themes and Variations? Front Syst Neurosci, 2017. 11: p. 80.

27. Ding, J.B., et al., Thalamic gating of corticostriatal signaling by cholinergic interneurons. Neuron, 2010. 67(2): p. 294–307.

28. Smith, Y., et al., The thalamostriatal system: a highly specific network of the basal ganglia circuitry. Trends Neurosci, 2004. 27(9): p. 520–7.

29. Lapper, S.R. and J.P. Bolam, Input from the frontal cortex and the parafascicular nucleus to cholinergic interneurons in the dorsal striatum of the rat. Neuroscience, 1992. 51(3): p. 533–45.

30. Matsumoto, N., et al., Neurons in the thalamic CM-Pf complex supply striatal neurons with information about behaviorally significant sensory events. J Neurophysiol, 2001. 85(2): p. 960–76.

31. Deffains, M. and H. Bergman, Striatal cholinergic interneurons and cortico-striatal synaptic plasticity in health and disease. Mov Disord, 2015. 30(8): p. 1014–25.

32. J. N. Reynolds RA, P.D.D., S. D. Fisher, M. J. Oswald, J. R. Wickens, Y. F. Zhang Coincidence of cholinergic pauses, dopaminergic activation and depolarisation of spiny projection neurons drives synaptic plasticity in the striatum. Nat Commun, 2022.

33. Borden, P.M., et al., A fast genetically encoded fluorescent sensor for faithful in vivo acetylcholine detection in mice, fish, worms and flies. bioRxiv, 2020.

34. Pancani, T., et al., Cholinergic deficits selectively boost cortical intratelencephalic control of striatum in male Huntington’s disease model mice. Nat Commun, 2023. 14(1): p. 1398.

35. Dong, C., et al., Fluorescence Imaging of Neural Activity, Neurochemical Dynamics, and Drug-Specific Receptor Conformation with Genetically Encoded Sensors. Annu Rev Neurosci, 2022. 45: p. 273–294.

36. Guo, Q., et al., Whole-brain mapping of inputs to projection neurons and cholinergic interneurons in the dorsal striatum. PLoS One, 2015. 10(4): p. e0123381.

37. Ma, T., et al., Chronic alcohol drinking persistently suppresses thalamostriatal excitation of cholinergic neurons to impair cognitive flexibility. J Clin Invest, 2022. 132(4).

38. Ma, T., et al., Bidirectional and long-lasting control of alcohol-seeking behavior by corticostriatal LTP and LTD. Nat Neurosci, 2018. 21(3): p. 373–383.

39. Gangal, H., et al., Drug reinforcement impairs cognitive flexibility by inhibiting striatal cholinergic neurons. Nat Commun, 2023. 14(1): p. 3886.

40. Becchi, S., et al., Cognitive effects of thalamostriatal degeneration are ameliorated by normalizing striatal cholinergic activity. Sci Adv, 2023. 9(25): p. eade8247.

41. Wu, Y.W., et al., Input- and cell-type-specific endocannabinoid-dependent LTD in the striatum. Cell Rep, 2015. 10(1): p. 75–87.

42. Diaz-Hernandez, E., et al., The Thalamostriatal Projections Contribute to the Initiation and Execution of a Sequence of Movements. Neuron, 2018. 100(3): p. 739–752.

43. Cheng, Y., et al., Distinct synaptic strengthening of the striatal direct and indirect pathways drives alcohol consumption. Biological Psychiatry, 2017. 81(11): p. 918–929.

44. Fleming, W., et al., Cholinergic interneurons mediate cocaine extinction in male mice through plasticity across medium spiny neuron subtypes. Cell Rep, 2022. 39(9): p. 110874.

45. Atack, J.R., et al., Molecular forms of acetylcholinesterase and butyrylcholinesterase in the aged human central nervous system. J Neurochem, 1986. 47(1): p. 263–77.

46. Shen, W., et al., M4 muscarinic receptor signaling ameliorates striatal plasticity deficits in models of L-DOPA-induced dyskinesia. Neuron, 2015. 88(4): p. 762–73.

47. Jeon, J., et al., A subpopulation of neuronal M4 muscarinic acetylcholine receptors plays a critical role in modulating dopamine-dependent behaviors. J Neurosci, 2010. 30(6): p. 2396–405.

48. Roberts-Wolfe, D., et al., Drug Refraining and Seeking Potentiate Synapses on Distinct Populations of Accumbens Medium Spiny Neurons. J Neurosci, 2018. 38(32): p. 7100–7107.

49. Bobadilla, A.C., et al., Cocaine and sucrose rewards recruit different seeking ensembles in the nucleus accumbens core. Mol Psychiatry, 2020. 25(12): p. 3150–3163.

50. Sullivan, M.A., H. Chen, and H. Morikawa, Recurrent inhibitory network among striatal cholinergic interneurons. J Neurosci, 2008. 28(35): p. 8682–90.

51. Koos, T. and J.M. Tepper, Dual cholinergic control of fast-spiking interneurons in the neostriatum. J Neurosci, 2002. 22(2): p. 529–35.

52. English, D.F., et al., GABAergic circuits mediate the reinforcement-related signals of striatal cholinergic interneurons. Nat Neurosci, 2011. 15(1): p. 123–30.

53. Faust, T.W., et al., Neostriatal GABAergic Interneurons Mediate Cholinergic Inhibition of Spiny Projection Neurons. J Neurosci, 2016. 36(36): p. 9505–11.

54. Bamford, N.S., R.M. Wightman, and D. Sulzer, Dopamine’s effects on corticostriatal synapses during reward-based behaviors. Neuron, 2018. 97(3): p. 494–510.

55. Chang, C.Y., et al., Brief optogenetic inhibition of dopamine neurons mimics endogenous negative reward prediction errors. Nat Neurosci, 2016. 19(1): p. 111–6.

56. van Zessen, R., et al., Cue and Reward Evoked Dopamine Activity Is Necessary for Maintaining Learned Pavlovian Associations. Journal of Neuroscience, 2021. 41(23): p. 5004–5014.

57. Salinas-Hernandez, X.I., et al., Dopamine neurons drive fear extinction learning by signaling the omission of expected aversive outcomes. Elife, 2018. 7.

58. Salinas-Hernandez, X.I., et al., Functional architecture of dopamine neurons driving fear extinction learning. Neuron, 2023.

59. Quinn, D.M., Acetylcholinesterase: enzyme structure, reaction dynamics, and virtual transition states. Chemical Reviews, 1987. 87(5): p. 955–79.

60. Ravel, S., E. Legallet, and P. Apicella, Responses of tonically active neurons in the monkey striatum discriminate between motivationally opposing stimuli. J Neurosci, 2003. 23(24): p. 8489–97.

61. Mineur, Y.S. and M.R. Picciotto, The role of acetylcholine in negative encoding bias: Too much of a good thing? Eur J Neurosci, 2021. 53(1): p. 114–125.

62. DiFiglia, M., Synaptic organization of cholinergic neurons in the monkey neostriatum. J Comp Neurol, 1987. 255(2): p. 245–58.

63. Gonzales, K.K., et al., GABAergic inputs from direct and indirect striatal projection neurons onto cholinergic interneurons in the primate putamen. J Comp Neurol, 2013. 521(11): p. 2502–22.

64. Martone, M.E., et al., Ultrastructural examination of enkephalin and substance P input to cholinergic neurons within the rat neostriatum. Brain Res, 1992. 594(2): p. 253–62.

65. Schulz, J.M., M.J. Oswald, and J.N. Reynolds, Visual-induced excitation leads to firing pauses in striatal cholinergic interneurons. J Neurosci, 2011. 31(31): p. 11133–43.

66. Kharkwal, G., et al., Parkinsonism driven by antipsychotics originates from dopaminergic control of striatal cholinergic interneurons. Neuron, 2016. 91(1): p. 67–78.

67. Sanchez, G., et al., Reduction of an afterhyperpolarization current increases excitability in striatal cholinergic interneurons in rat parkinsonism. J Neurosci, 2011. 31(17): p. 6553–64.

68. Zhang, Y.F., J.N.J. Reynolds, and S.J. Cragg, Pauses in cholinergic interneuron activity are driven by excitatory input and delayed rectification, with dopamine modulation. Neuron, 2018. 98(5): p. 918–925 e3.

69. Apicella, P., et al., The role of striatal tonically active neurons in reward prediction error signaling during instrumental task performance. J Neurosci, 2011. 31(4): p. 1507–15.

70. Ravel, S., E. Legallet, and P. Apicella, Tonically active neurons in the monkey striatum do not preferentially respond to appetitive stimuli. Exp Brain Res, 1999. 128(4): p. 531–4.

71. Carnicella, S., D. Ron, and S. Barak, Intermittent ethanol access schedule in rats as a preclinical model of alcohol abuse. Alcohol, 2014. 48(3): p. 243–52.

72. Wei, X., et al., Dopamine D1 or D2 receptor-expressing neurons in the central nervous system. Addict Biol, 2018. 23(2): p. 569–584.

73. Ma, T., et al., Bidirectional and long-lasting control of alcohol-seeking behavior by corticostriatal LTP and LTD. Nat Neurosci, 2018. 21(3): p. 373–383.

74. Hellard, E.R., et al., Optogenetic control of alcohol-seeking behavior via the dorsomedial striatal circuit. Neuropharmacology, 2019. 155: p. 89–97.

75. Ma, T., et al., Alcohol induces input-specific aberrant synaptic plasticity in the rat dorsomedial striatum. Neuropharmacology, 2017. 123: p. 46–54.

76. Cheng, Y., et al., Prenatal exposure to alcohol induces functional and structural plasticity in dopamine D1 receptor-expressing neurons of the dorsomedial striatum. Alcohol Clin Exp Res, 2018. 42(8): p. 1493–1502.

77. Huang, C.C.Y., et al., Stroke triggers nigrostriatal plasticity and increases alcohol consumption in rats. Sci Rep, 2017. 7(1): p. 2501.

78. Wang, J., et al., Ethanol-mediated long-lasting adaptations of the NR2B-containing NMDA receptors in the dorsomedial striatum. Channels (Austin), 2011. 5(4): p. 205–209.

79. Lu, J., et al., Alcohol intake enhances glutamatergic transmission from D2 receptor-expressing afferents onto D1 receptor-expressing medium spiny neurons in the dorsomedial striatum. Neuropsychopharmacology, 2019. 44(6): p. 1123–1131.

80. Cui, G., et al., Concurrent activation of striatal direct and indirect pathways during action initiation. Nature, 2013. 494(7436): p. 238–42.

81. Wang, J., et al., Alcohol elicits functional and structural plasticity selectively in dopamine D1 receptor-expressing neurons of the dorsomedial striatum. J Neurosci, 2015. 35(33): p. 11634–43.

82. Franklin, K.B.J. and G. Paxinos, The mouse brain in stereotaxic coordinates. 3 ed. 2007, San Diego: Academic Press.

83. Paxinos, G. and C. Watson, The Rat Brain in Stereotaxic Coordinates. 2008, New York: Academic Press.

84. Lu, J., et al., Whole-brain mapping of direct inputs to dopamine D1 and D2 receptor-expressing medium spiny neurons in the posterior dorsomedial striatum. eNeuro, 2021. 8(1): p. 0348-20.2020.

