## Supplementary Figures 1-7 for "Dynamic Responses of Striatal Cholinergic Interneurons Control the Extinction and Updating of Goal-Directed Learning"

### Supplemental Figures

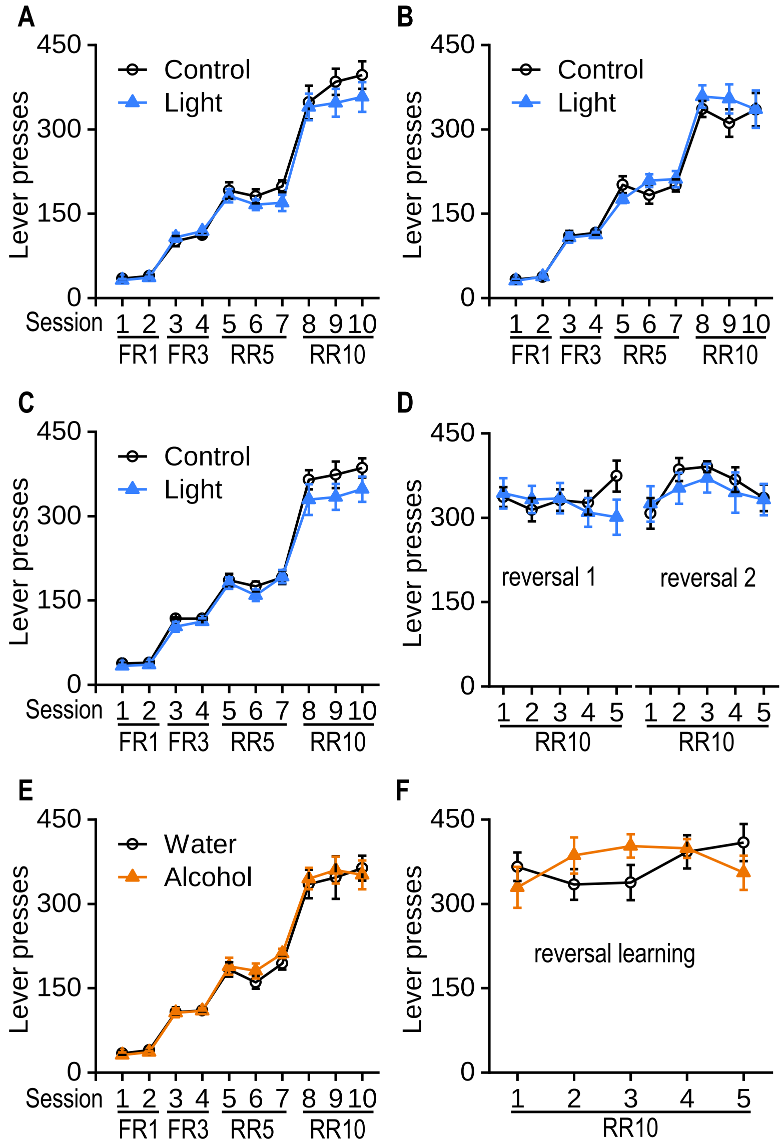


**Supplemental Figure 1. Learning curves of instrumental conditioning.** Rats were trained in the operant chambers to press a lever for rewards, progressing from fixed ratio (FR) protocols FR1 and FR3 to random ratio (RR) protocols RR5 and RR10. Reversal learning used the RR10 protocol. A. The learning curve of animals is in Figure 1. There was a main effect of session ($F_{(1,9)}$ = 156.99, $p$ < 0.001) but no group effect ($F_{(1,216)}$ = 1.45, $p$ = 0.24). n = 13 rats (Control) and 13 rats (Light). **B.** The learning curve of animals is in

Figure 2. There was a main effect of session ($F_{(1,9)}$ = 120.76, $p < 0.001$) but no group effect ($F_{(1,135)}$ = 0.54, $p = 0.48$). n = 8 rats (Control) and 9 rats (Light). **C.** The initial learning curve of animals is in Figure 3. There was a main effect of session ($F_{(1,9)}$ = 217.29, $p <$ 0.001) but no group effect ($F_{(1,261)}$ = 2.40, $p = 0.13$). **D.** The reverse learning curve of animals is in Figure 3. There was no main group effect ($F_{(1,261)}$ = 0.25, $p = 0.62$). n = 16 rats (Control) and 15 rats (Light). **E.** The initial learning curve of animals is in Figure 5. There was a main effect of session ($F_{(1,9)}$ = 144.96, $p < 0.001$) but no group effect ($F_{(1,171)}$ = 0.14, $p = 0.71$). **F.** The reversal learning curve of animals is in Figure 5. There was no main group effect ($F_{(1,76)}$ = 0.27, $p = 0.61$). n = 11 rats (Water) and 10 rats (Alcohol).

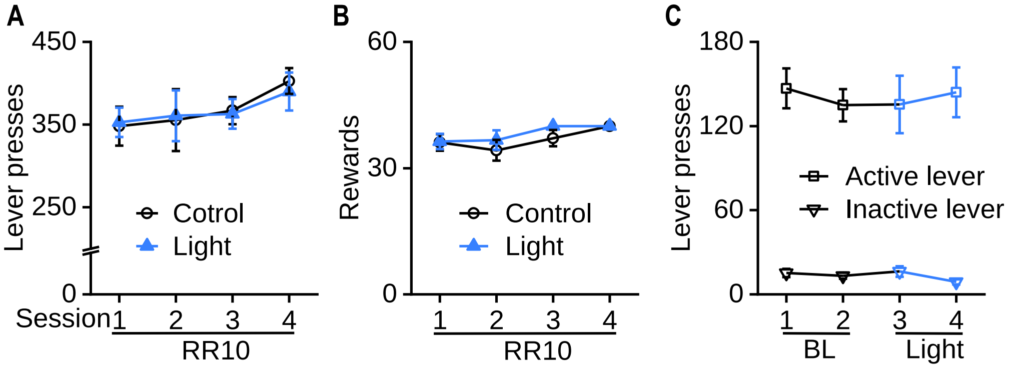


**Supplemental Figure 2. Optical stimulation of CINs with rewards delivery. A.** Rats were trained in the operant chambers to press a lever for rewards using the RR10 protocol. Lever presses triggered reward and synchronized light stimulation (590 nm) time-locked to reward delivery. Light stimulation comprised 5 repetitions of 20 Hz light bursts (10 pulses, 5 ms per pulse) with a 0.8-second interval. There was no main group effect ($F_{(1,48)}$ = 0.007, $p$ = 0.94). **B.** The earned rewards during training were not significantly different between the two groups ($F_{(1,48)}$ = 0.90, $p$ = 0.36). n = 9 rats (Control) and 9 rats (Light). **C.** Rats were trained in the operant chambers to press the active lever for a reward using the FR3 protocol. Once animals reached two stable baseline (BL) sessions performance, light stimulation (590 nm) time-locked to reward delivery was administrated for the next two sessions. Light stimulation comprised 5 repetitions of 20 Hz light bursts (10 pulses, 5 ms per pulse) with a 0.8-second interval. There was no main session effect ($F_{(1,3)}$ = 0.23, $p$ = 0.88). n = 7 rats.

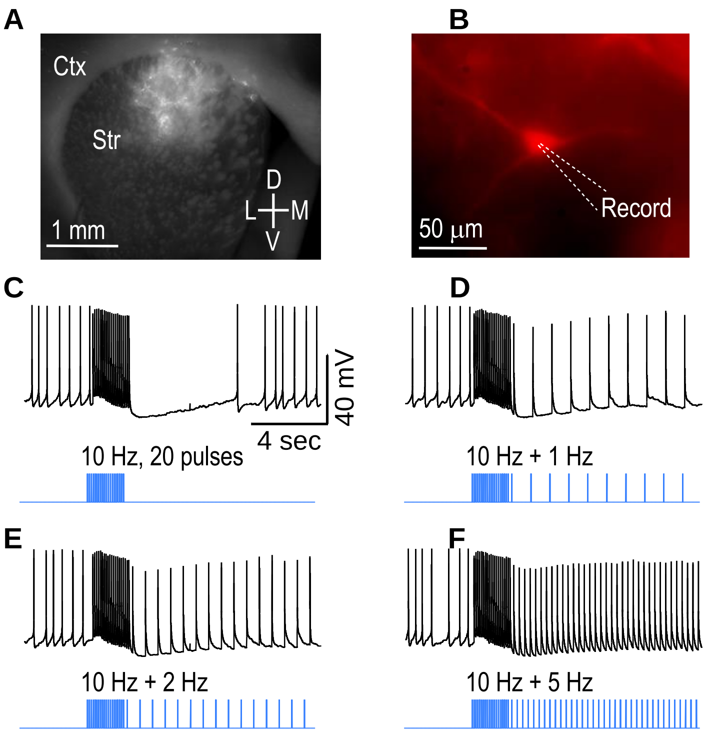


**Supplemental Figure 3. Disrupting pause with continuous optical stimulations in current-clamp Recording. A.** The expression of AAV-FLEX-ChrimsonR-tdTomato in the DMS of ChAT-Cre rat. **B.** Recording of tdTomato expressed CIN using whole-cell patch clamp. **C.** Optical stimulation (590 nm, 10 Hz, 20 pulses) induced a burst-pause firing in the CIN. Disruption of the pause in CIN firing through continuous optical stimulation with different frequencies: **D.** 1 Hz, **E.** 2 Hz, **F.** 5 Hz.

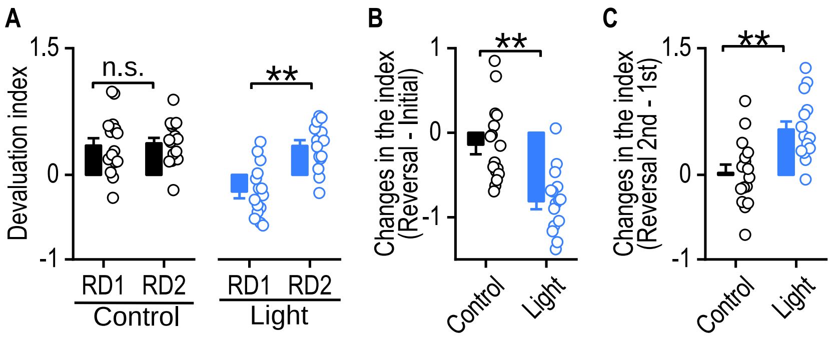


**Supplemental Figure 4: Further analysis of devaluation index data of Figure 5. A.** The devaluation index was not significantly different between reversal devaluation 1 (RD1) and reversal devaluation 2 (RD2) in the control group; Unpaired *t* test, $t_{(30)} = -0.24$, $p = 0.81$, n.s., not significant; the devaluation index was significantly higher in the RD2 than the RD1 for the light group, indicating a recovered reversal learning; Unpaired *t* test, $t_{(28)} = -5.13$, $^{**}p < 0.01$. **B.** The changes in devaluation index between RD1 and initial devaluation were significantly lower in the light group than the control group, indicating an impaired reversal learning; Unpaired *t* test, $t_{(29)} = 4.47$, $^{**}p < 0.01$. **C.** The changes of devaluation index between RD2 and the RD1 was significantly higher in the light group than the control group, indicating a recovered reversal learning; Unpaired *t* test, $t_{(29)} = -3.73$, $^{**}p < 0.01$. n = 16 rats (Control) and 15 rats (Light).

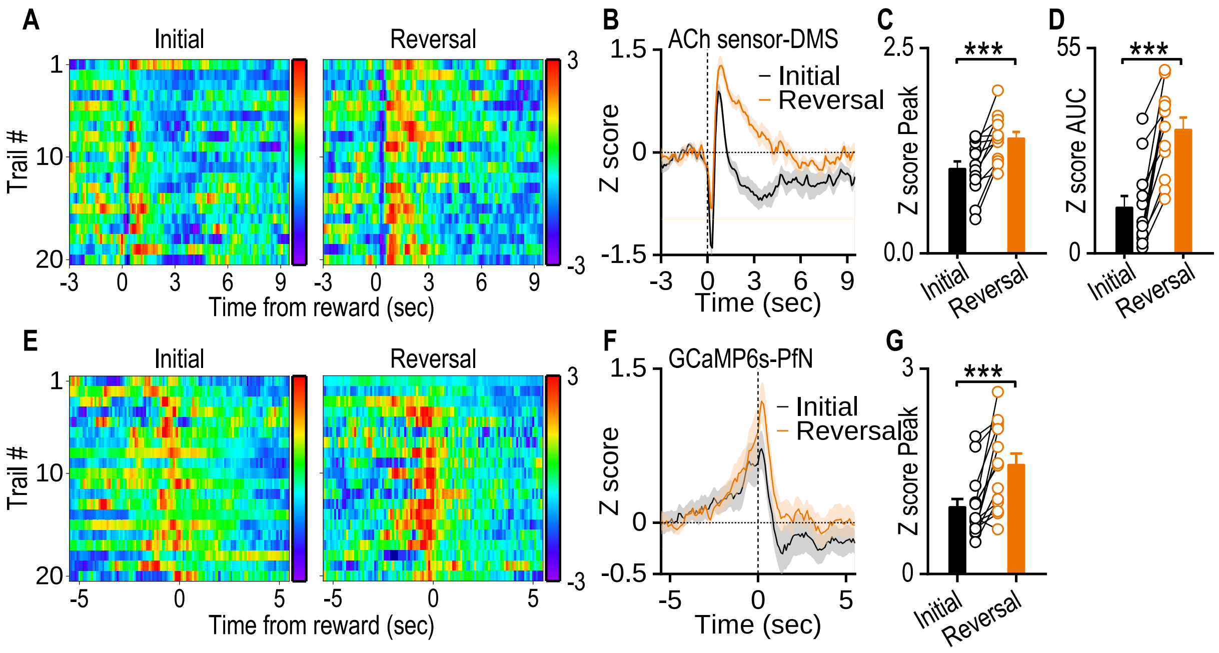


**Supplemental Figure 5. Increased ACh release and enhanced PfN activity during reversal learning in Long-Evans rats.**

**A.** *In vivo* measurements of ACh signals in the DMS. The heat map showed ACh signals during the first reversal session, which consisted of initial learning in the first half session and reversal learning in the second half. **B.** Representative traces of ACh signals during the first reversal session. **C.** Summary data quantifying the peak ACh sensor fluorescence signals; Paired *t* test, $t_{(11)} = -4.71$, ***$p < 0.001$. **D.** Summary data quantifying the area under the curve (AUC) of ACh sensor fluorescence signals; Paired *t* test, $t_{(11)} = -6.61$, ***$p < 0.001$. **E.** *In vivo* measurements of GCaMP signals in the PfN. The heat map showed GCaMP signals during the first reversal session, which consisted of initial learning in the first half session and reversal learning in the second half. **F.** Representative traces of GCaMP signals during the first reversal session. **G.** Summary data quantifying the peak

GCaMP sensor fluorescence signals, Paired *t* test, $t_{(13)} = -4.41$, ***$p < 0.001$. Paired *t* test, n = 12 sessions from 6 rats for (C, D), n = 14 sessions from 7 rats for (G).

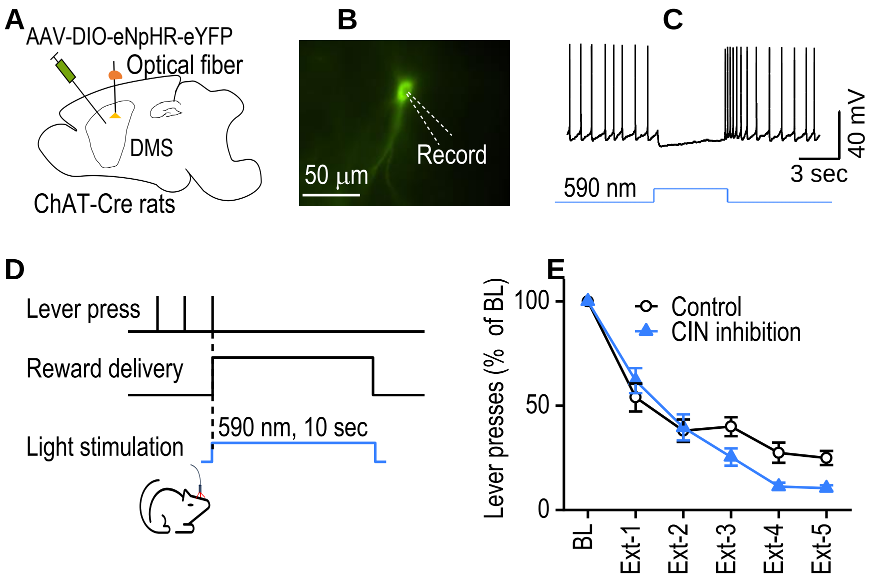


**Supplemental Figure 6. CIN inhibition does not slow down the extinction of goal-directed learning. A.** Schematic of viral injection and optical fiber implantation. ChAT-Cre rats received a bilateral infusion of AAV-DIO-eNpHR-eYFP, and optical fibers were bilaterally implanted into the DMS. Rats then underwent instrumental training to receive rewards by pressing levers. **B.** Recording of eYFP expressed CIN using whole-cell patch clamp. **C.** Optical stimulation (590 nm, 5 sec) inhibited the firing of CIN. **D.** Optical stimulation protocol employed during extinction training. Lever presses triggered both reward and synchronized light stimulation (590 nm) time-locked to reward delivery. Light was continuously given for 10 seconds during the reward delivery period. Actual rewards were omitted during extinction. **E.** Lever presses during extinction training were not significantly different between the light-stimulation and control groups. The data were normalized to their baseline lever presses. There was no main group effect ($F_{(1,88)} = 2.68$, $p = 0.12$). n = 12 rats (Control) and 12 rats (Light).

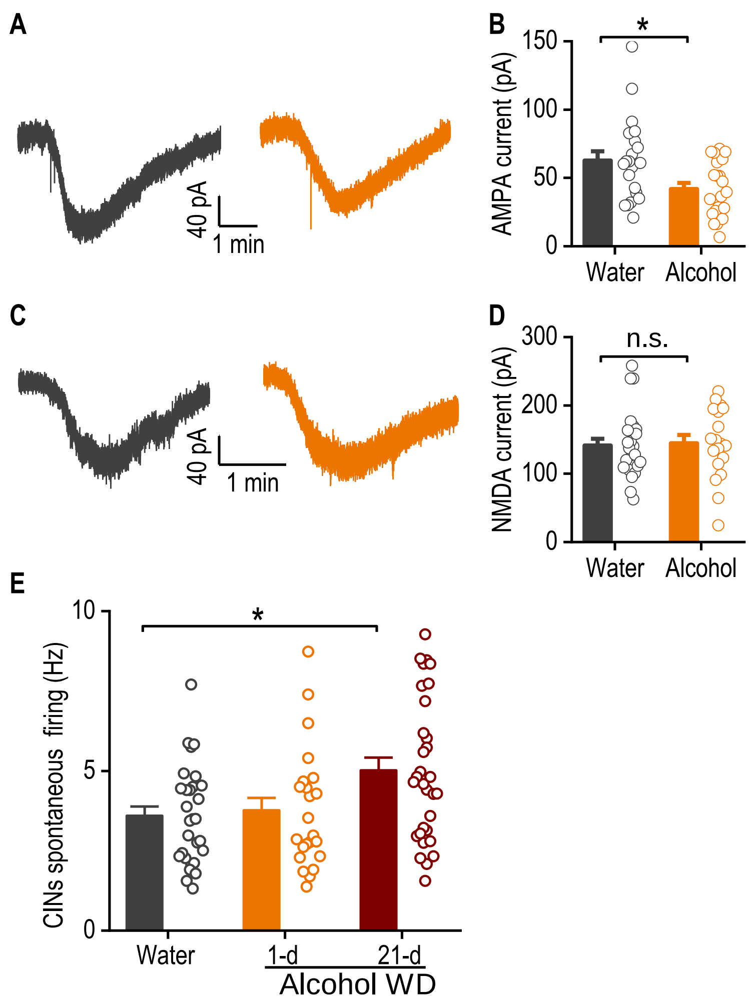


**Supplemental Figure 7. Chronic alcohol intake reduces AMPA-induced currents in DMS CINs. A.** Sample traces of bath application of AMPA (10 µM, 15 seconds) induced currents of CIN in water (left) and alcohol (right) drinking animals. **B.** Summary data quantifying the peak currents. Unpaired *t* test, $*p < 0.05$, n = 21 neurons from 5 mice (Water 21/5) and (Alcohol 21/4). **C.** Sample traces of bath application of NMDA (30 µM, 30 seconds) induced currents of CIN in water (left) and alcohol (right) drinking animals. NMDA-induced currents were recorded in a magnesium-free external solution. **D.** Summary data quantifying the peak currents. Unpaired *t* test, $p = 0.82$, n = (Water 24/5) and (Alcohol 19/4). **E.** Spontaneous firing rates of CINs in the indicated groups; one-way

ANOVA $F_{(2,80)} = 4.79$, $p = 0.01$, $*p < 0.05$ vs. water group by Tukey *post hoc* test; n = (Water 29/5), (Alcohol WD1-d 23/5), and (Alcohol WD 21-d 31/5).
